# Towards Autonomous Intra-cortical Brain Machine Interfaces: Applying Bandit Algorithms for Online Reinforcement Learning

**DOI:** 10.1101/2020.01.08.899641

**Authors:** Shoeb Shaikh, Rosa So, Tafadzwa Sibindi, Camilo Libedinsky, Arindam Basu

## Abstract

This paper presents application of Banditron - an online reinforcement learning algorithm (RL) in a discrete state intra-cortical Brain Machine Interface (iBMI) setting. We have analyzed two datasets from non-human primates (NHPs) - NHP A and NHP B each performing a 4-option discrete control task over a total of 8 days. Results show average improvements of ≈ 15%, 6% in NHP A and 15%, 21% in NHP B over state of the art algorithms - Hebbian Reinforcement Learning (HRL) and Attention Gated Reinforcement Learning (AGREL) respectively. Apart from yielding a superior decoding performance, Banditron is also the most computationally friendly as it requires two orders of magnitude less multiply-and-accumulate operations than HRL and AGREL. Furthermore, Banditron provides average improvements of at least 40%, 15% in NHPs A, B respectively compared to popularly employed supervised methods - LDA, SVM across test days. These results pave the way towards an alternate paradigm of temporally robust hardware friendly reinforcement learning based iBMIs.

## I. INTRODUCTION

With as many as 1 in 50 people suffering from paralysis worldwide [1], Intra-cortical Brain Machine Interfaces (iBMIs) have a great potential in restoring semblance of a normal life for these embattled souls. iBMIs literally transform thought into action. The input to this system is neural signals recorded from the surface of the motor cortex to drive effectors such as cursors [2], prosthetic limbs [3], wheelchairs [4] and so on.

Current iBMI systems involve time-consuming daily calibration procedures [5], thereby lending inconvenience to the end-users. These procedures are based on supervised learning methods which require explicit measurement of effector kinematic variables. The overall systems reported so far often involve a neuro-engineer to be present to perform these procedures and get the system up and running for the end-user.

An alternative to supervised learning methods is *reinforcement learning* (RL) wherein a model is updated on the fly in response to a simple scalar reward [6]–[9] (see Fig. 1). Briefly put, an RL agent takes *states* and *rewards* as inputs and yields an output *action* at a given time-step. *States* correspond to the binned spike firing rates sensed from micro-electrode arrays whereas *rewards* are scalar values obtained as a result of *action* taken at the previous time-step. The notion of learning involves maximizing the score of total reward. This approach has the potential to tackle the following existing problems,

- Daily calibration routines are not required as the algorithm is updated online while being in use
- Explicit measurement of effector kinematic variables is not required and a scalar reward signal is enough
- Bringing the above two points in fruition has the potential to take the neuro-engineer out of the loop

**Fig. 1.**
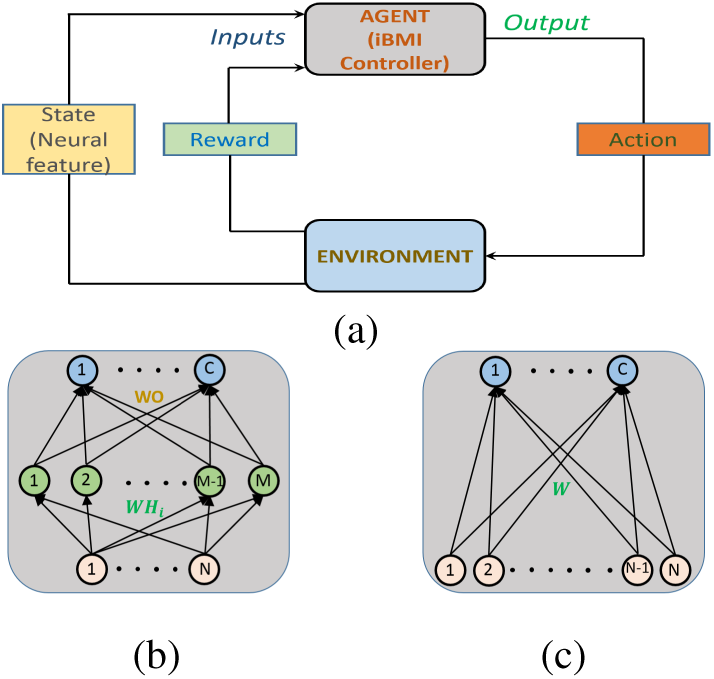
(a) Block level representation of Reinforcement Learning in context of iBMIs. The iBMI controller emits an output action based on inputs - Neural states and rewards. For an Agent (iBMI Controller) with *N* input electrodes and *C* target labels, (b) represents architectural scheme of two layer neural network based RL algorithms - AGREL and HRL, whereas (c) depicts single layer neural network based RL algorithm - Banditron.

Encouraging RL based results have been reported in [6] where authors have applied Q-learning in a discrete state iBMI involving rats trained to control a prosthetic arm to choose among two targets. However, Q-learning suffers from generalization difficulty owing to the curse of dimensionality problem resulting from the large neural state-action space. To address this issue, authors in [8] have proposed Attention Gated Reinforcement Learning (AGREL) [10] as an alternative and have found it to be convincingly better than Q-learning in a cursor based center-out task involving one NHP. In AGREL [8], authors resort to computation of the instantaneous reward based on the distance of the cursor from the target. [7], [11] on the other hand report a Hebbian Reinforcement learning (HRL) algorithm based on a discrete binary reward having values of only +1 or −1 in a two target NHP based iBMI setup. However, state of the art RL algorithms - AGREL and HRL suffer from generalization problems as they employ neural networks that are susceptible to the local optimum problem [9]. To tackle this problem, we propose applying Banditron - an online prediction algorithm used popularly in sponsored advertising and recommender systems [12].

## II. REINFORCEMENT LEARNING ALGORITHMS

### A. State of art RL algorithms applied in iBMIs

#### 1) AGREL

It involves a three-layer neural network that maps the input neural state to action space. Output of the *j*^*th*^ hidden layer node is given as,

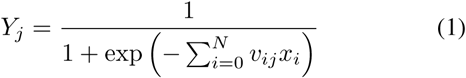

where *N* is the number of input electrodes, *x*_*i*_ is the firing rate appearing at the *i*^*th*^ electrode, *v*_*ij*_ corresponds to the input layer weights and *x*_0_ = 1 to account for input weight bias.

Stochastic softmax rule is employed by AGREL to select a winning neuron amidst *C* output neurons. Value of *k*^*th*^ output neuron, *Z*_*k*_, and its associated probability is given as,

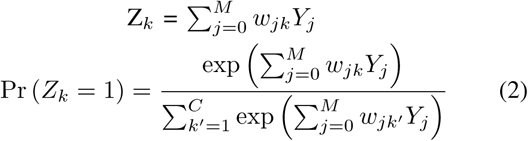

where *w*_*jk*_ represents the output layer weights, *M* is the number of hidden nodes.

Instantaneous reward is given as,

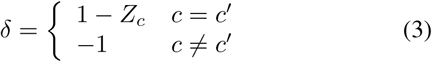

where *c* and *c′* are predicted and true actions respectively.

AGREL resorts to an instantaneous reward-based expansive function, *f* (*δ*), to expedite learning,

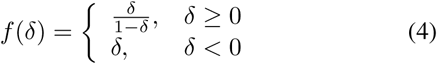

Randomly initialized weights are updated at every time-step as follows,

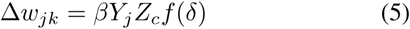

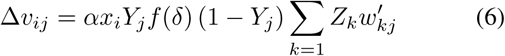

where *α* and *β* are learning rates.

#### 2) HRL

It employs a three-layer neural network similar to AGREL with a different way of approaching learning. The hidden node output is given as,

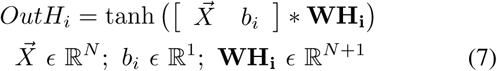

where 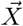 represents the input feature vector, *b*_*i*_ represents a bias term and **WH**_**i**_ stands for input weights.

The output nodes have action values given as,

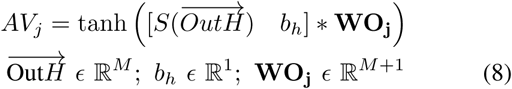

where *S*() represents the sign function, *b*_*h*_ represents a bias term and **WO**_**j**_ represent the output weights. Highest valued output node is selected as the final output. Feedback reward signal, *f*, is given as,

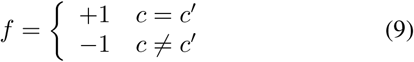

where *c* and *c′* are predicted and true actions respectively.

Randomly initialized weights are updated at every time-step as follows,

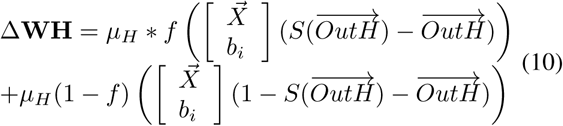

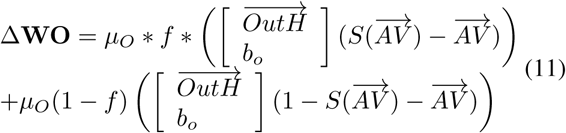

where 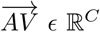 is the action value output vector.

### B. Proposed RL algorithm - Banditron

At time-step *t* = 1, Banditron weight matrix starts out as ***W*** ^**1**^ = **0** ∈ ℝ^*C*×*N*^. Subsequently, at *t* = 1, 2, …, *T*, with inputs, *x*_*t*_ *ϵ* ℝ^*N*^, Banditron employs current weight matrix ***W*** ^*t*^ to yield,

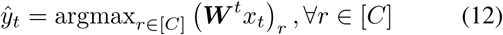

where [*C*] = 1, …, *c. γ* is a parameter that sets the exploration-exploitation tradeoff. Accordingly, Banditron leaves itself for exploration with probability 1 − *γ* and uniformly picks a random label from [*C*] as 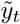 which is the final predicted label. Feedback signal is 1 if actual output, *y*_*t*_ equals final predicted value, 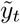 and 0 otherwise. The weight matrix is updated as,

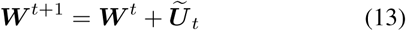

where 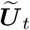 represents the update matrix. True label (*y*_*t*_) is not revealed, however, we implicitly obtain full information when, 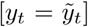. Thus, as seen in Fig. 2, update matrix 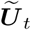 is written as a function of randomized 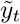.

**Fig. 2.**
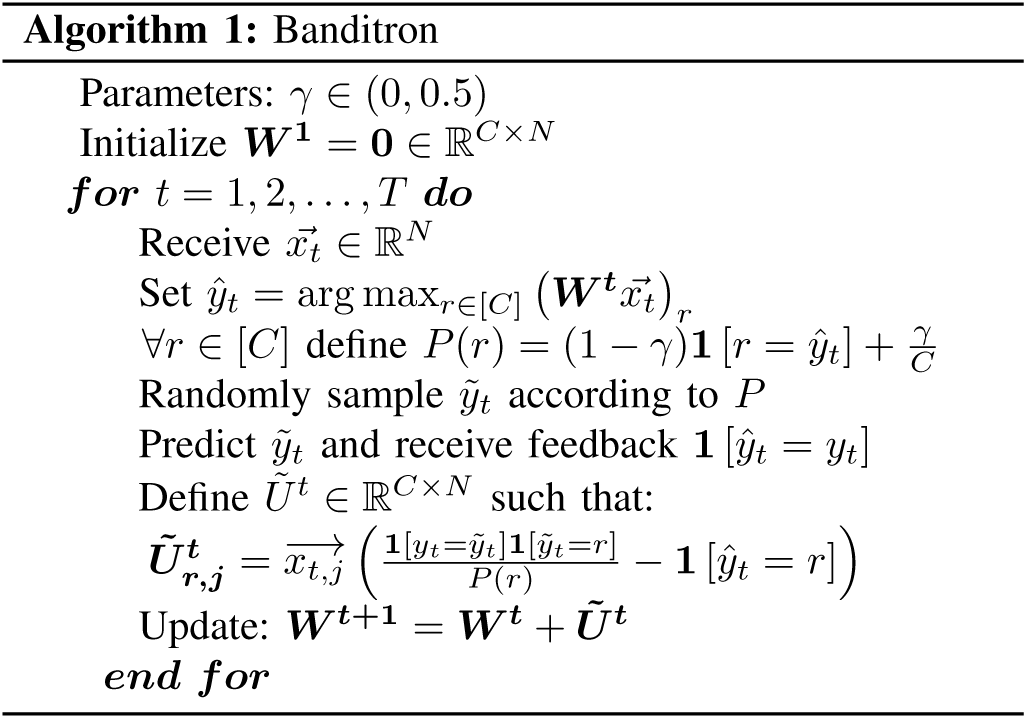
Banditron pseudocode [12]

Banditron is a modification of simple linear perceptron and hence is capable of learning only linearly separable patterns [12]. However, non-linear methods have shown superior performance over linear methods in iBMIs [13], [14]. Thus, with this evidence we introduce non-linearity in the form of obtaining feature vector - *f*_*t*_ *ϵ* ℝ^*M*^ as non-linear random projection (RP) of *x*_*t*_ given as,

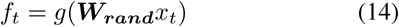

where ***W***_***rand***_ *ϵ* ℝ^*N* ×*M*^ is the fixed random projection weights, *g*(.) corresponds to the activation function which in our case is sigmoid. Learning proceeds in the same manner as described in Fig. 2 with 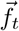 serving as input instead of 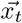. We henceforth refer to this variant of Banditron with input 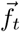 as *Banditron* − *RP*.

## III. METHODS

### A. Behavioural task and data acquisition

NHP A was trained to control a robotic wheelchair through a 3-directional joystick as shown in Fig. 3. The experiment consisted of four tasks - a) Moving forward by 2 m, b) turning left by 90°, c) turning right by 90° and d) staying still for 5 seconds. The experiment was carried out in the form of trials wherein a movement related trial was considered successful if NHP A reached the target within 15 seconds. NHP B on the other hand was trained to control a similar joystick to manipulate a cursor on screen. A trial in this experiment corresponded to a square-shaped target being shown at top, right or left positions relative to the centre of the screen. A trial was considered successful if the NHP managed to move the cursor from the centre of the screen to the target area and stay in it for 1.5 seconds within a total elapsed time of 13 seconds.

**Fig. 3.**
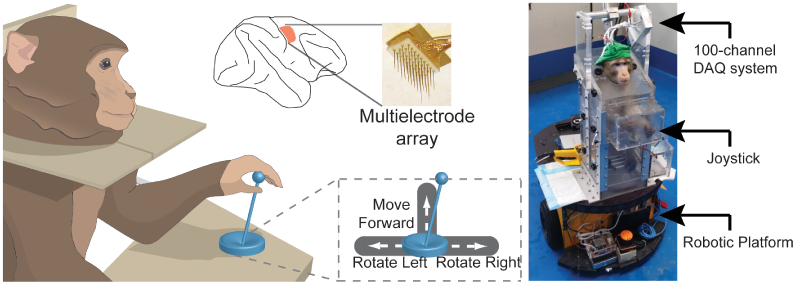
NHP A was trained to move a robotic wheelchair through a 3-directional joystick [4] (CC-BY license).

Both NHPs A and B had a total 64 electrodes sensing raw neural data from the motor cortex area. Threshold crossings [15] at each of the input electrodes were used to compute the resultant firing rates in time-bins of 500 ms. Firing rates were used as inputs to the iBMI system.

### B. Analysis Methodology

We have used a total of 8 days of experimental data with 3 recorded sessions for analysis in both NHPs. The iBMI decoder takes binned firing rates as an input and outputs one among the following discrete actions - *Right, Forward, Left* or *Stop* every 100 ms. State of the art methods used in iBMIs are supervised learning based methods. Hence, we present comparison of popular supervised learning methods in BMIs - Linear Discriminant Analysis (LDA) and Support Vector Machines (SVM) [16] against RL methods - AGREL, HRL and Banditron. We present comparison of two variants of supervised models - *fixed* and *retrained. fixed* supervised models are trained on the first session of *Day* 1 whereas *retrained* supervised models are trained on the first session of the respective test day across both NHPs. Do note that the RL methods do not require a separate training session and are trained on the fly across the *test* sessions in both NHPs with a binary reward at the end of every prediction. Reported RL methods are initialized from scratch every single day. The test sessions compared are consistent across all the reported methods.

## IV. RESULTS

### A. Hyperparameter tuning

Algorithms - SVM, HRL, AGREL, Banditron and Banditron-RP have hyper-parameters that require tuning. For SVM we have tuned the hyperparameters based on the training set, while using bayesian optimization function *bayesopt* provided in MATLAB2019. HRL, AGREL, Banditron and Banditron-RP are RL algorithms and thus do not employ an explicit training set to tune weights. However, they have hyper-parameters to be tuned and we have chosen these based on *Day* 1′*s* first session data in both NHPs.

### B. Decoder Results

**Fig. 4.**
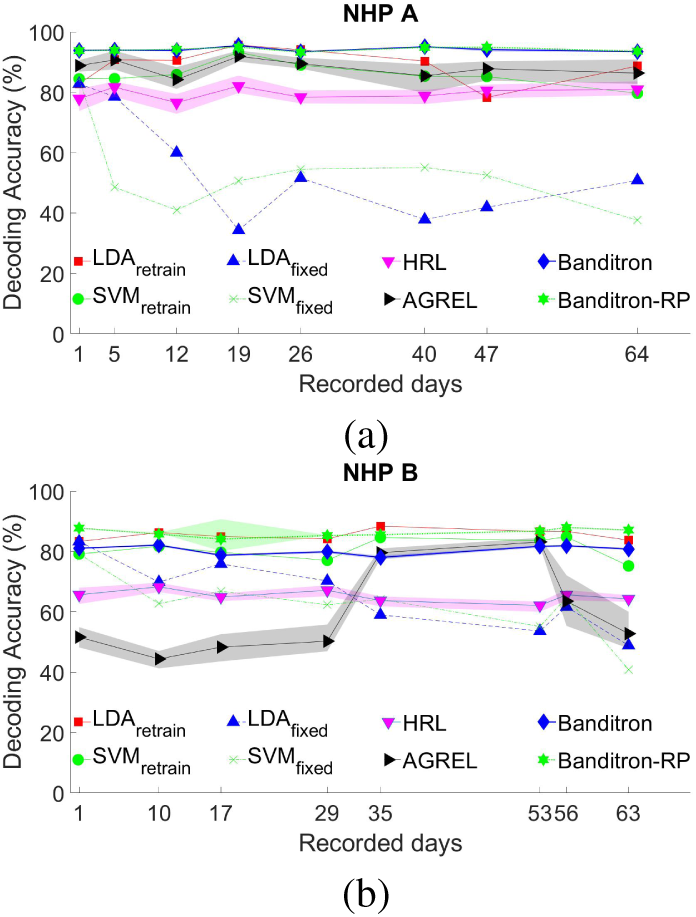
Decoded results over 8 days for different models in (a) NHP A and (b) NHP B. Shaded bands represent variance over 20 random instantiations in RL algorithms.

Banditron yields an average improvement of 6.01%, 14.5% in NHP A and 20.98%, 14.97% in NHP B over AGREL, HRL respectively. Furthermore, Banditron compares favorably with retrained supervised models while yielding average improvement of at least ≈ 5% in NHP A while reaching up to 95% level of performance. Do note that the comparison is not an even one as the *training* methodology differs. Retrained supervised algorithms are fixed after training on session 1 of a given day with full label information. RL algorithms on the other hand do not have an explicit training session but are updated with a binary feedback throughout the *test* set. Supervised learning based results, thus, are presented not for direct comparison but for establishing state of the art context for the presented RL methods.

The non-linear variant of Banditron, Banditron-RP performs as good as Banditron in NHP A and yields improvement of ≈ 6% in NHP B. Statistical comparison of Banditron against HRL and AGREL yield p-values of 0.0078, 0.0078 in NHP A and 0.0078, 0.0391 in NHP B respectively for a Wilcoxon signed-rank test. This shows that the improvements afforded by Banditron are statistically significant (*p <* 0.05) over HRL, AGREL.

### C. Computational Complexity Comparison

We consider an iBMI system with *N* inputs and *C* option discrete control for computation. The number of multiply-and-accumulate operations (MACs) required during prediction for single layer classifiers such as LDA, Banditron can be given as,

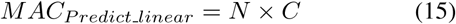

For single hidden layer neural network based approaches such as HRL, AGREL with *M* hidden nodes and a bias term in both input-hidden (*WH*_*i*_) and hidden-output weight (*WO*) matrices, the number of MACs required during prediction is given as,

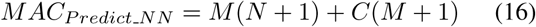

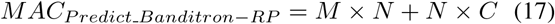

We have ignored the number of MACs required to implement the activation function for the sake of simplicity.

RL algorithms are recursively updated and MACs expended by single-layer Banditron and multi-layer neural network systems - AGREL, HRL and Banditron-RP can be formulated as below,

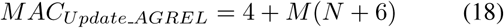

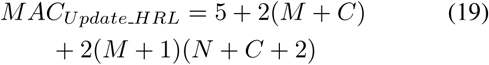

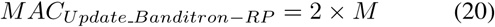

For a single layer Banditron based network, number of MACs required per update is given as,

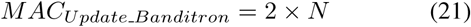

Do note that state of the art supervised schemes such as LDA, SVM are trained at the beginning of the day/session and are held fixed without being updated iteratively.

Consider a case of a *N* = 64 input electrode iBMI system driving a 4-option (*C* = 4) discrete control similar to reported experiments in this paper. For neural networks based approaches HRL and AGREL, let us assume number of hidden nodes to be *M* = 80. With these assumptions, we can arrive at the Table. I comparing number of MACs and memory requirement across algorithms. Note that we have considered a linear version of SVM in the below reported table and we consider weights to be 16-bits across all algorithms.

**TABLE I.** COMPUTATIONAL COMPLEXITY ACROSS DIFFERENT MODELS

| Algorithms | MACs<br>update | MACs<br>prediction | Memory<br>(kB) |
| --- | --- | --- | --- |
| LDA, SVM | - | 256 | $N \times C \times W_{res} = 0.5$ |
| AGREL | 5604 | 5524 | $[M(N+1) + C(M+1)]W_{res} = 10.8$ |
| HRL | 11513 | 5524 | $[M(N+1) + C(M+1)]W_{res} = 10.8$ |
| Banditron | 64 | 256 | $N \times C \times W_{res} = 0.5$ |
| Banditron-RP | 160 | 5440 | $[M(N+C)]W_{res} = 10.6$ |

MAC = multiply and accumulate, Number of input channels, $N = 64$ , Number of hidden layer nodes, $M = 80$ , Number of outputs, $C = 4$ , Weight resolution( $W_{res}$ ) = 16bits.

## V. DISCUSSION

### A. Comparison

It seems counter-intuitive that a single layer network architecture - Banditron outperforms two layer neural network architectures - AGREL, HRL. The reason for Banditron’s superior performance is its amenability to learn in an online setting [12]. Multi-layer neural network based approaches like HRL, AGREL suffer from convergence issues when trained in an online manner. They are typically trained in batches over multiple epochs [8].

### B. Reward Signal

Currently, we are using an ideal binary feedback signal processed in a post-hoc fashion. Research has shown this signal to be present in real-time in biological signal sources in the brain such as nucleus accumbens [17], primary motor cortex [18], [19]. Alternatively, one can also derive these signals from other signal sources such as EEG [20], ECoG [21]. Certainly, more work is needed to a) fetch a reliable source of reward signal and b) study the reward signal’s impact on the iBMI system with regards to its frequency and accuracy.

## VI. CONCLUSION AND FUTURE WORK

We have presented Banditron as an alternative computationally friendly technique to existing state of the art RL-iBMI techniques with statistically significant improvements across two NHPs. Furthermore, validation of Banditron has also been presented against supervised learning techniques.

We will explore teasing out reward signal from a biological source such that future iBMI systems are fully autonomous and co-adaptive in the true sense. Applying RL-algorithms in a continuous state iBMI setup is a goal we wish to undertake in future.

## Notes

** This work was supported through grant RG87/16 by MOE, Singapore

## REFERENCES

[1] Armour, Courtney-Long, Fox, Fredine, et al., “Prevalence and causes of paralysis-united states, 2013.” American journal of public health, vol. 106 10, pp. 1855–7, 2016.

[2] Pandarinath, Nuyujukian, Blabe, Sorice, et al., “High performance communication by people with paralysis using an intracortical brain-computer interface,” eLife, p. e18554, 2017.

[3] Collinger, Wodlinger, Downey, Wang, et al., “High-performance neuroprosthetic control by an individual with tetraplegia.” Lancet (London, England), no. 9866, pp. 557–64, 2013.

[4] Libedinsky, So, Xu, Kyar, et al., “Independent mobility achieved through a wireless brain-machine interface,” PLoS ONE, vol. 11, no. 11, pp. 1–13, 2016.

[5] Brandman, Burkhart, Kelemen, Franco, et al., “Robust Closed-Loop Control of a Cursor in a Person with Tetraplegia using Gaussian Process Regression,” Neural Computation, vol. 30, no. 11, pp. 2986–3008, 2018.

[6] DiGiovanna, Mahmoudi, Fortes, Principe, et al., “Coadaptive Brain–Machine Interface via Reinforcement Learning,” IEEE Transactions on Biomedical Engineering, vol. 56, no. 1, pp. 54–64, Jan. 2009.

[7] Pohlmeyer, Mahmoudi, Geng, Prins, et al., “Using Reinforcement Learning to Provide Stable Brain-Machine Interface Control Despite Neural Input Reorganization,” PLoS ONE, vol. 9, no. 1, 2014.

[8] Wang, Wang, Xu, Zhang, et al., “Neural Control of a Tracking Task via Attention-Gated Reinforcement Learning for Brain-Machine Interfaces,” IEEE Transactions on Neural Systems and Rehabilitation Engineering, vol. 23, no. 3, pp. 458–467, May 2015.

[9] Zhang, Member, Libedinsky, So, et al., “Clustering Neural Patterns in Kernel Reinforcement Learning Assists Fast Brain Control in Brain-Machine Interfaces,” IEEE Transactions on Neural Systems and Rehabilitation Engineering, vol. 4320, no. c, pp. 1–10, 2019.

[10] Roelfsema and Ooyen, “Attention-gated reinforcement learning of internal representations for classification,” Neural Computation, vol. 17, no. 10, pp. 2176–2214, 2005.

[11] Mahmoudi, Pohlmeyer, Prins, Geng, et al., “Towards autonomous neuroprosthetic control using Hebbian reinforcement learning,” Journal of Neural Engineering, vol. 10, no. 6, p. 066005, Dec. 2013.

[12] Kakade, Shalev-Shwartz, and Tewari, “Efficient bandit algorithms for online multiclass prediction,” Proceedings of the 25th International Conference on Machine Learning, pp. 440–447, 2008.

[13] Sussillo, Stavisky, Kao, Ryu, et al., “Making brain-machine interfaces robust to future neural variability,” Nature Communications, pp. 1–12.

[14] Shaikh, So, Sibindi, Libedinsky, et al., “Towards Intelligent Intracortical BMI (i2BMI): Low-power Neuromorphic Decoders that out-perform Kalman Filters,” IEEE Transactions on Biomedical Circuits and Systems, pp. 1–1, 2019.

[15] Trautmann, Stavisky, Lahiri, Ames, et al., “Accurate estimation of neural population dynamics without spike sorting,” Neuron, vol. 103, no. 2, pp. 292 – 308.e4, 2019.

[16] Lotte, Bougrain, Cichocki, Clerc, et al., “A review of classification algorithms for EEG-based brain–computer interfaces: a 10 year update,” Journal of Neural Engineering, vol. 15, no. 3, p. 031005, apr 2018.

[17] Prins, Sanchez, and Prasad, “Feedback for reinforcement learning based brain-machine interfaces using confidence metrics,” Journal of Neural Engineering, vol. 14, no. 3, 2017.

[18] Marsh, Tarigoppula, Chen, and Francis, “Toward an Autonomous Brain Machine Interface: Integrating Sensorimotor Reward Modulation and Reinforcement Learning,” Journal of Neuroscience, vol. 35, no. 19, pp. 7374–7387, 2015.

[19] Benyamini, Nason, Chestek, and Zacksenhouse, “Neural Correlates of error processing during grasping with invasive brain-machine interfaces,” in 2019 9th International IEEE/EMBS Conference on Neural Engineering (NER). IEEE, Mar., pp. 215–218.

[20] Kreilinger, Neuper, and Müller-Putz, “Error potential detection during continuous movement of an artificial arm controlled by brain– computer interface,” Medical & biological engineering & computing, vol. 50, no. 3, pp. 223–230, 2012.

[21] Milekovic, Ball, Schulze-Bonhage, Aertsen, et al., “Error-related electrocorticographic activity in humans during continuous movements,” Journal of neural engineering, vol. 9, no. 2, p. 026007, 2012.

